# PRENATAL STRESS MODIFIES SPATIAL COGNITION IN MICE: EFFECTS OF AGE AND SEX

**DOI:** 10.1101/2024.02.23.581828

**Authors:** M McCarthy, TG Buck, MS Sodhi

**Affiliations:** Department of Molecular Pharmacology and Neuroscience, Loyola University Chicago, Maywood, IL 60153

## Abstract

Prenatal stress is linked with neuropathology of the cortical-hippocampal circuit, due to abnormal brain development that leads to long term neurological deficits in offspring, in both rodents and in humans with psychiatric disorders. Conflicting reports exist of the effects of prenatal stress in C57BL/6J mice, which is an inbred strain that is the most frequently used for neurobehavioral research. We now present comprehensive analyses of the effects of prenatal stress on spatial cognition and related behaviors in this inbred strain, in males and females at young adult and aged adult stages. The prenatal stress exposure was conducted by exposing pregnant mice to restraint stress three times daily from day 7 to day 18 of gestation. We tested the effects of prenatal stress on cognitive behavior using a battery of behavioral assays including the open-field test, the light-dark box, Morris water maze spatial acquisition and spatial reversal testing, in addition to assays of social interaction and social memory. We compared the behavioral phenotype of male and female offspring in young adult (10-20 weeks) and older adult ages (60- 70 weeks). Data reveal that male but not female mice had reduced body weight throughout the lifespan after exposure to prenatal stress. Young prenatally stressed adult females showed greater organization of tracking behavior in spatial acquisition tests than other groups. Prenatal stress improved reversal learning in aged females relative to non-stressed aged females. We detected decreased extinction of the spatial acquisition memory in prenatally stressed young adult mice of both sexes. Overall results indicate that prenatal stress may protect females from detrimental effects of age on function of the hippocampus. Our data support the finding that C57BL/6J are a relatively resilient strain of mouse that may be useful for investigating the differences between resilience and susceptibility to stress. Genomic differences between the C57BL/6J strain relative to Swiss Webster strain, which is more susceptible to prenatal stress are discussed. Future studies of prenatal stress in the C57BL/6J mouse strain should focus on the identification of molecular pathways that underlie resilience to prenatal stress, to identify novel targets for drug development.

## Introduction

Human epidemiological studies have indicated prenatal stress is associated with offspring depression, anxiety, autism spectrum disorders and schizophrenia (Abbott et al., 2018). For example, in a study of the *Dutch Hunger Winter* in the Netherlands during World War II, female offspring of women exposed to severe food deprivation were at an increased risk of developing schizophrenia (Susser & Lin, 1992), while, offspring of women consuming less than 1,000 kcal per day during the second and third trimester, were at increased risk of major depression (Brown et al., 2000). Another study showed that rates of schizophrenia were increased in patients born in 1960 and 1961 in mothers exposed to the Chinese famine of 1969-1961 (St Clair et al., 2005). Stress is not only linked to physiological stressors; when bereavement of a first-degree relative during pregnancy is used as a measure of stress, there is an increased relative risk of schizophrenia (Khashan et al., 2008). In contrast, Class et al. (2014), a large retrospective cohort study in Sweden, found no association between schizophrenia and bereavement but did find positive associations with suicide, autism spectrum disorders, and attention deficit hyperactivity disorder.

Despite the heterogeneous, and sometimes contradictory, outcomes, prenatal stress has continued to be used in rodent studies with exposure to various stressors, immune challenges, infections, and malnutrition (Weinstock, 2017). This has resulted in a wide range of outcomes even within the same inbred strains of rodents, as in human studies. These differences between strains have been attributed to phenotypes that recapitulate some features of anxiety (Miyagawa et al., 2011), depression (Behan et al., 2011; Clarke et al., 2019; Sierksma et al., 2013), schizophrenias (Dong et al., 2015; Matrisciano et al., 2018; Matrisciano et al., 2013; Matrisciano et al., 2012), and autism spectrum disorders (Dunn et al., 2024). The strains in each study varied such that prenatally stressed C57BL/6J mice exhibited behaviors associated with risk aversion while prenatally stressed Swiss Webster mice were more vulnerable to deficits in social behaviors. At a microscopic level, prenatal stress has been shown to affect glutamate receptors (Matrisciano et al., 2012; Orlando et al., 2014), GABAegic proteins(Matrisciano et al., 2018; Matrisciano et al., 2012), serotonin (Miyagawa et al., 2011), the HPA axis (Mueller & Bale, 2008; Salomon et al., 2011; Son et al., 2006), BDNF and DNMT (Dong et al., 2015; Matrisciano et al., 2012), RNA editing (Bristow et al., 2021), long-term potentiation (Son et al., 2006), and total glial cell count (Behan et al., 2011).

In the current study, we sought to identify behavioral deficits resulting from prenatal restraint stress in young and old adult C57BL/6J mice of both sexes. This inbred strain is widely used as a background to test the neurophysiological and behavioral effects of genetic modifications but has a specific phenotype that might bias studies of stress. We performed a comprehensive battery of behaviors to assess the effect of prenatal stress in this commonly used inbred strain. We conducted the open field test to analyze locomotor activity, immobile episodes, and risk tolerance behaviors. We performed the light-dark box test as a secondary measure of risk aversion. We used the the Morris water maze assay to analyze spatial acquisition learning and spatial reversal learning. These tests were performed in both young adult mice (10-20 weeks) and aged adult mice (60-70 weeks) of both sexes. We also measured social behavior in young adults.

Data reveal that male but not female mice had reduced body weight throughout the lifespan after exposure to prenatal stress. Although there were no differences associated with prenatal stress in the open field, light/dark box, or social behavior, we detected several age and sex differences in the effects of prenatal stress on spatial acquisition and spatial reversal memory. Overall results indicate that prenatal stress may protect females from detrimental effects of age on function of the hippocampus. Our data support the finding that C57BL/6J are a relatively resilient strain of mouse that may be useful for investigating the differences between resilience and susceptibility to stress. Genomic variation in the C57BL/6J strain may influence its resilient phenotype.

## Methods

### Animals and prenatal stress procedure

Male and female C57BL/6J mice (#000664) were purchased from The Jackson Laboratory (Bar Harbor, ME), and were maintained in same-sex groups, in standard caging, in a temperature- and humidity-controlled room (22°C, 40-55% humidity), on a 12-hour light-dark cycle. Mice were given access to food and water *ad libitum*. The animal facility was AAALAC-approved, and all procedures were performed in accordance with protocols approved by the Institutional Animal Care and Use Committee at Loyola University Chicago.

Breeder pairs were time-mated at 4 months of age. The females were then individually housed and randomly assigned to one of two groups: control, n = 6; or prenatal restraint stress (PRS), n = 8. The day after pairing was considered gestation day 0. Control dams were left undisturbed throughout gestation, while PRS dams were subjected to repeated episodes of restraint stress. From gestation day 7 to gestation day 18, PRS dams were exposed to restraint stress. Dams were removed from their cages and placed into a transparent restraining tube, under a bright light, for 45 minutes, 3 times per day. The offspring from 6 PRS dams and 3 control dams were used as subjects in this study, the remaining dams did not produce litters.

Offspring from control and PRS dams were weaned on postnatal day 21, housed in same-stress same-sex pairs, and maintained on a 12-hour *reverse* light-dark cycle. Due to a small number of successful litters from the control dams, and an imbalance in the sex of the offspring in the litters, only female subjects were used from our control breeders. Male control subjects were purchased from Jackson Laboratories, Behavioral comparisons (Morris water maze, open field test, etc.) between the purchased mice and age matched controls bred in-house showed no significant differences (Supplementary Figure 1). The male control mice were age-matched to the male PRS subjects and delivered to our facility at 5 weeks of age and allowed to acclimatize for 3-4 weeks before testing. A total of 40 subjects were used in this study, 10 subjects per group (PRS/male, PRS/female, control/male, control/female).

### Behavior testing

These tests were performed in both young adult mice (10-20 weeks) and aged adult mice (60-70 weeks) of both sexes unless otherwise specified. Subjects began behavioral testing at 2- 4 months of age. Mice were acclimated to a 12-hour reverse light dark cycle more than 2 weeks before testing. All tests were conducted in the same room, separate from the room where the subjects were housed, during the dark phase of the animals’ light cycle. The behavioral testing battery included: 1) the *open field locomotor activity* test, to assess general activity level; 2) the *light-dark box test*; 3) the *Morris water maze*, to test spatial learning and memory, and 4) social interaction and social novelty to test sociability with other mice as well as social memory.

Two experimenters tested the mice and were blind to treatment conditions throughout testing. All experiments were conducted using ANY-maze behavior tracking software (Stoelting Co., Wood Dale, IL). Experiments, *except social interaction and novelty,* were repeated using the same offspring mice approximately 1 year after the first behavioral tests began.

### Open field test

These tests were performed in both young adult mice (10-20 weeks) and aged adult mice (60-70 weeks) of both sexes. Each subject was given one 30-minute open field locomotor activity test session. The subject was placed into the center of a square, opaque, plastic chamber, measuring 50 cm (length) x 50 cm (width) x 30 cm (height). Room lighting was dim and indirect. Testing order was counterbalanced across experimental groups, and the chamber was cleaned with 95% ethanol before and after each session, to remove scent cues. The tracking software recorded total distance travelled and time spent immobile. In addition, the software recorded the time spent in each of two zones: the *center zone*, a 25 cm x 25 cm square area centered in the 50 cm x 50 cm chamber; and the *periphery zone*, between the walls of the chamber and the center zone.

### Light-dark box test

These tests were performed in both young adult mice (10-20 weeks) and aged adult mice (60-70 weeks) of both sexes. Each mouse subject was given one 10-minute light-dark box test session. The subject was placed into the center of a square, transparent, plastic chamber, under a bright light. This open chamber measured 30 cm (length) x 30 cm (width) x 30 cm (height) and was connected to another chamber via a small opening. The second chamber had black walls and a black ceiling and measured 15 cm (length) x 30 cm (width) x 30 cm (height). The testing order was counterbalanced across experimental groups, and the chamber was cleaned with 95% ethanol before and after each session, to remove scent cues. The tracking software recorded the time spent in the light and dark chambers, and the number of entries into the dark chamber.

### Morris water maze (MWM)

These tests were performed in both young adult mice (10-20 weeks) and aged adult mice (60-70 weeks) of both sexes. The water maze was a circular pool (170 cm in diameter) made of opaque, white plastic. A circular escape platform (12 cm in diameter), made of clear plastic, was submerged 1 cm below the water surface (San Diego Instruments, San Diego, CA). The water temperature was maintained at ambient air temperature and was measured to be 19-21 °C. Room lighting was dim and indirect. The maze was surrounded by high-contrast, distal cues (approximately 90 cm away from the pool).

The tracking software divided the maze into four equal quadrants with two perpendicular lines, the end of each line demarcated four cardinal points: North (N), South (S), East (E), and West (W). The escape platform was always positioned in the approximate center of one of the four quadrants. The swim path length and latency to find the escape platform were recorded.

The Morris water maze protocol used was adopted from Vorhees and Williams (2006). Briefly, mice began with the **cued learning t**ask as a control procedure to ensure all subjects had the necessary functions to complete the task. Each subject was given four trials per day for three days. A bright neon tennis ball was positioned 12 cm above the water and above the escape platform. The subject was placed in the pool facing the wall at one of the cardinal locations. Each trial lasted 90 seconds or until the subject reached the platform. If, after 90 seconds, the subject did not find the platform, they would be guided to the platform. Each subject was allowed to remain on the platform for 15 seconds before being returned to the home cage.

The second task was **spatial acquisition**. The marker (tennis ball) above the platform was removed. Each subject was assigned an island location before starting, which was counterbalanced across all groups. Each trial lasted 90 seconds or until the subject found the platform. If the subject did not find the platform within 90 seconds, they were guided to the platform. All subjects were tested for four trials per day for 5 days. On the sixth day, a probe trial was conducted. In this probe trial, the platform was removed, and the subject was allowed to freely explore the water maze for a total of 90 seconds. After the 90 seconds elapsed the subject was removed from the water maze, dried with a dry towel, and returned to their home cage.

Lastly, the subjects underwent **spatial reversal learning** two days after the spatial acquisition probe trial. The location of the hidden platform was changed from southwest island to northeast, or from northeast island locations to southwest for each of the subjects. The same protocol as spatial acquisition was followed for each of the trials. All subjects were tested for 5 days with one probe trial one day after the last reversal trial.

**Trackplot categorization of Morris water maze probe trials** were made subjectively by viewing trackplots generated by Any-Maze. Trackplots were categorized into one of chaining, scanning, random, thigmotaxis, and focal search. Categorizations were defined according to Cooke et al. (2019). Chaining was defined as circular paths in all 4 quadrants. Scanning was defined as an average distance to the center of less than 60% of the radius with no apparent path. Random search was defined by no apparent path while spend more than 50% of the time one the outside of 60% of the radius. Thigmotaxis was defined as wall hugging at least 65% of the time. Focal search was defined by staying close to the target island. Trackplots generated by Any-Maze were blinded to the assessor. Direct path, directed search, and indirect search were not seen. The assessor performed 2 categorizations of each trackplot. Comparisons were made between the categorizations, and, if 2 straight categorizations agreed, the categorization was accepted. If there was disagreement, a third review was given as the final response (**Fig. 7A**). Categories were collapsed into two groups: organized (focal search or chaining) or disorganized (scanning, random, or thigmotaxis) for analysis. We used chi-square analysis to test for differences in search strategy due to age, sex and stress in both spatial acquisition probe trials and spatial reversal probe trials.

### Social Interaction and Social Memory

These tests were performed in only young adult mice (10-20 weeks) of both sexes but not in aged adult mice (60-70 weeks). Each mouse was tested on social interaction and social memory. Mice were tested using a three-chambered apparatus as previously described (Kaidanovich-Beilin et al., 2011). Briefly, after habituation to the apparatus, a stranger mouse was placed in one of the side chamber’s enclosed cages, while the other enclosed cage remained empty. Time spent in each chamber as well as a 2 cm greater radius than the enclosed cage surrounding each of the enclosed cages. Entries and exits from each zone were measured as well. A percentage of time of the whole was calculated as well as a *sociability index (Dunn et al., 2024)*, which was defined as entries into the x+2 cm diameter surrounding the stranger mouse’s enclosed cage divided by total entries into that same approach zone as well as the reciprocal zone surrounding the empty cage. Mice were allowed to freely explore the chamber for a total of 10 minutes.

After a brief rest period of 5 minutes, the test was repeated but the previously empty cage now contained a second stranger mouse, defined as the ‘novel stranger’ mouse. This second phase of the test measured **social memory**.

All stranger mice were age and gender matched to the mice being tested. The sides chosen for the stranger mice were counter-balanced between prenatally stressed and non-stressed groups. Female and male mice were tested on different days to avoid confounding scents from the chamber. In between each trial, the three-chamber apparatus as well as the enclosed cages were wiped clean with 95% ethanol.

### Statistical Analysis

All analyses were performed using GraphPad Prism (version 8.3; GraphPad Software, LLC, San Diego, CA). All measures were tested using ANOVA (if there were matched subjects) or mixed effects analysis (if there was subject drop out). Furthermore, weights were analyzed using 2 different time points – 6-10-week period resembling growth and development as well as an approximately 54-week time point resembling adult weights. Specific behavioral statistics are described in each of the corresponding sections above.

Furthermore, the Jax control NS male subjects were compared to a set of five NS control C57BL/6J mice bred at the Loyola facility, in the Morris Water Maze cued learning, spatial acquisition, and spatial reversal tasks. The mice were also compared using the probe trial data. There were no significant differences between the control bred mice and the Jax mice in the open field measurements or the Morris water maze cued learning, spatial acquisition, or spatial reversal (**Supplementary Figure 1**).

## Results

### Body weight

Male NS mice appeared to consistently outweigh PRS mice across the 54-weeks of measurements (Fig. 1A). Female mice in both NS and PRS groups appeared to overlap across the entire 54 weeks (Fig. 1B). Slight differences can be seen between the groups from 6-10 weeks (Fig. 1C). Lastly, differences in total body weight occurred at approximately 54 weeks (Fig. 1D).

**Figure 1:**
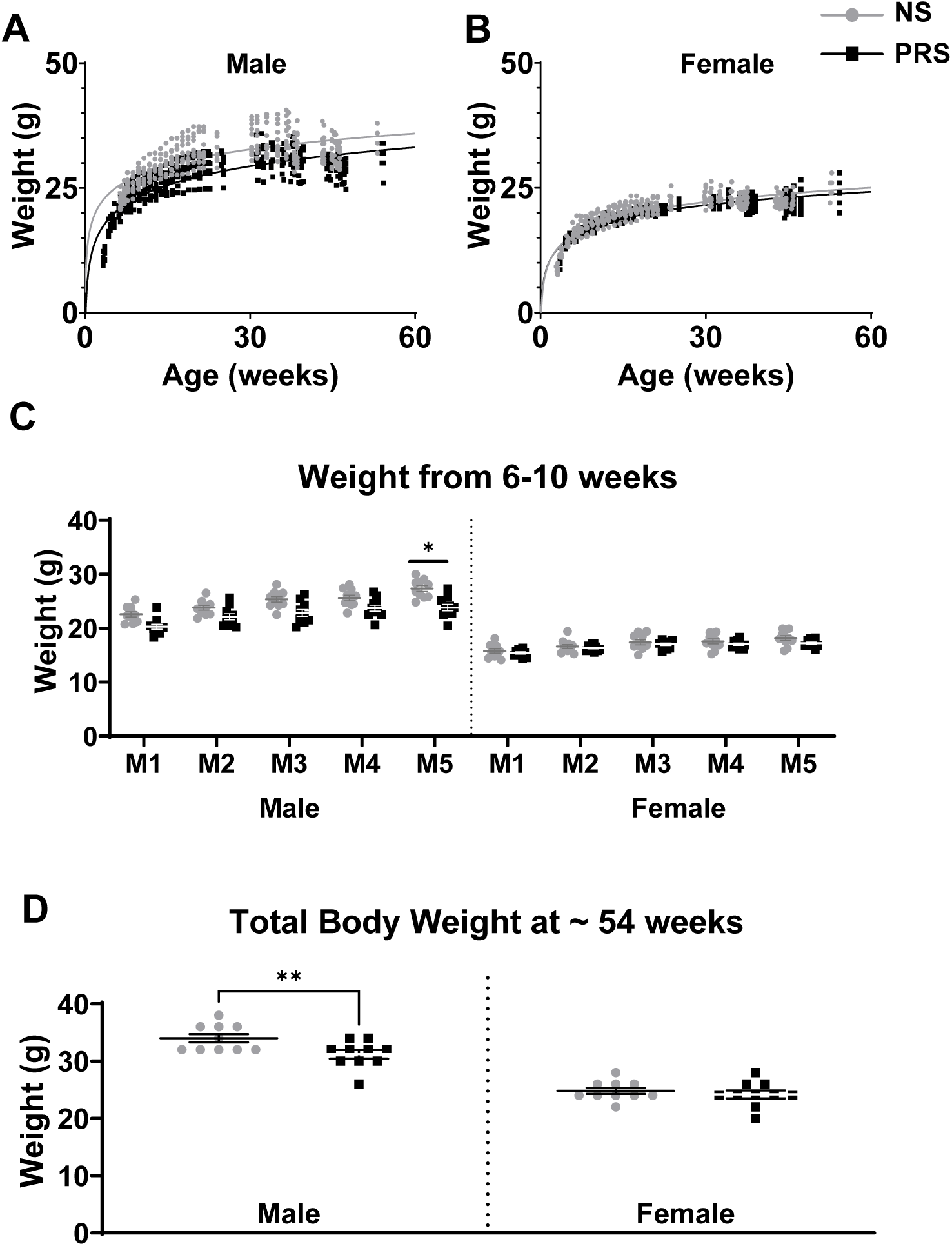
Body Weights of Mouse Cohort. **A, B.** The total body weight across 54 weeks of the experiment. Gray and black lines represent nonlinear semilog regression lines created for each of the groups of males and females. **C.** The weights from 6-10 weeks of development corresponding to young adulthood. M1-5 are each of the measurements that were taken as specific ages could not be used due to differences of specific days. M1 always starts at the first measurement after week 6. Exact ages were found to be insignificantly different when averaged, with PRS males and PRS females having the highest average age. **D.** The total body weight at the last measurement for each of the subjects in the experiment (approximately 54 weeks). *p<0.05, ** p < 0.01.

Analysis using ANOVA of body weight (g) at 6 through 10 weeks of age revealed a significant effect of sex (F_1,36_ = 259.7, *p* < 0.0001), a significant effect of stress (F_1,36_ = 11.20, *p* < 0.0001), and a significant interaction between sex and stress (F_1, 36_ = 4.872, p = 0.0338).

A two-way ANOVA of weight at the last time point (corresponding to approximately 54 weeks in each of the subjects) revealed a significant effect of stress (F_1,36_ = 6.237 p = 0.0172) and gender (F_1,36_=141.6, p < 0.0001). *Post-hoc* multiple comparisons using Fisher’s LSD exact test revealed a significant difference between NS Males and PRS Males (p=0.0062), but no significant difference between NS females and PRS females (p = 0.5370).

### Open field activity test

Data are summarized in Figure 2. Analysis of *<u>total distance</u>* using ANOVA (age x sex x stress) revealed a significant effect of age (F_1, 36_ = 10.16, p = 0.003) and sex (F_1, 36_ = 5.275, p = 0.027), but no significant effect of stress (F_1,36_ = 0.02772, p = 0.868). *Post-hoc* analysis using Sidak’s multiple comparisons revealed significantly lower activity in PRS males when young relative to activity when older (p = 0.0004). Moreover, young NS males had lower activity than young NS females (p = 0.0105).

**Figure 2:**
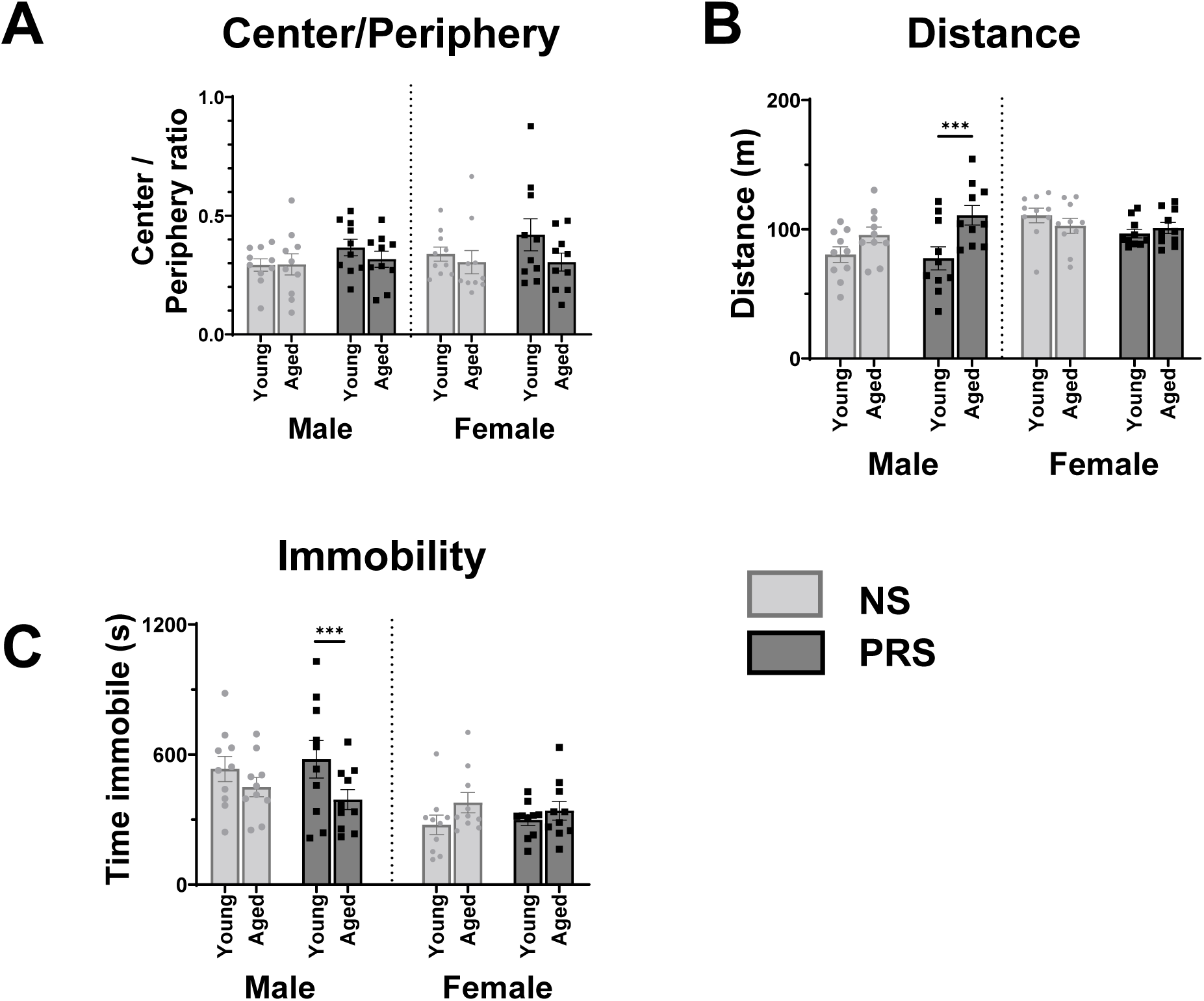
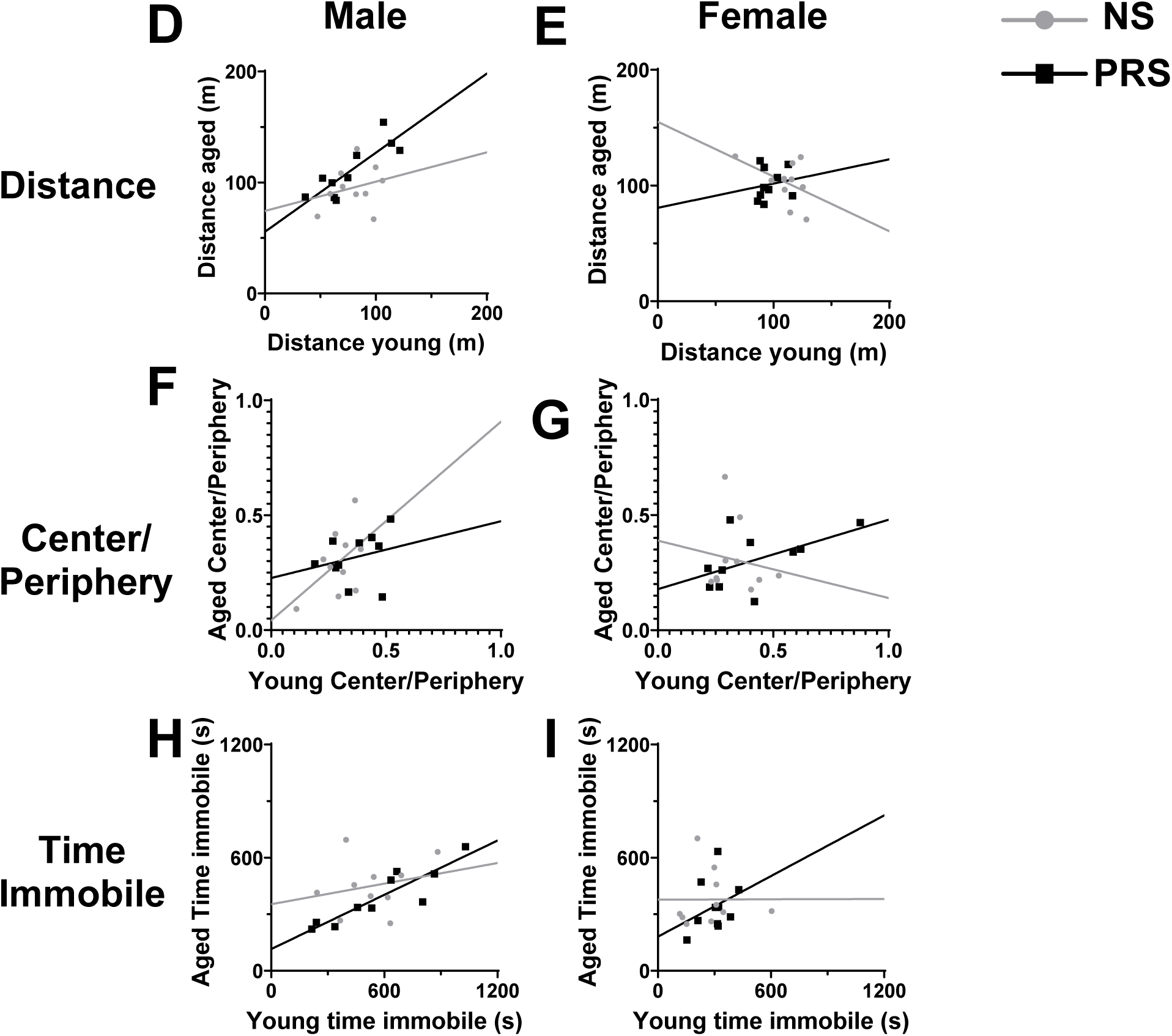
Open Field Testing. **A.** The time spent in the center of the field divided by the time in the periphery is an indicator of risk tolerance. Any measurement above 1.0 is suggestive of a preference for the center. **B.** The distance in meters for all the mice during the 900 second testing window is an indicator of locomotor activity. **C.** Time spent immobile is defined as > 1 second without movement, which is an indicator of fear. **D, E.** Correlations between young and aged time points for OFT distance in males and females, respectively. Multivariate ANOVA results showed a significant interaction between age and sex (F_1, 36_ = 13.81, p = 0.0007) and age and stress (F_1, 36_ = 4.718, p = 0.0365). **F, G.** Correlations between young and aged time points for OFT center/periphery in males and females, respectively. Multivariate ANOVA results showed no significant interactions of age and stress or age and sex. **H, I.** Correlations between young and aged time points for OFT time immobile for males and females, respectively. Multivariate analysis showed a significant interaction between age and gender (F_1, 36_ = 13.64, p = 0.0007) * p < 0.05, ** p < 0.01, *** p < 0.001, **** p < 0.0001

Analysis of time in the center divided by time in the periphery (center/periphery) using ANOVA (age x sex x stress) revealed no significant difference between any subgroup. Analysis of *<u>immobility</u>* using ANOVA revealed a significant effect of sex (F_1, 36_ = 14.24, p = 0.0006), but no significant effect of stress or age. The ANOVA did reveal a significant interaction between age and sex (F_1, 36_ = 13.64, p = 0.0007). *Post-hoc* analysis using Sidak’s multiple comparisons revealed significantly higher immobility in PRS males when younger (p = 0.0247). Young PRS males also had higher immobility relative to PRS females (p = 0.0036).

Simple linear regression analysis (Fig. 2D-I) compared each of the parameters of distance, center/periphery, and immobility between young adult and aged adult time points. The analysis revealed only significant deviation from 0 in PRS males in OFT distance (R^2^= 0.7148, p= 0.0021), and PRS males in immobility time (R^2^ = 0.8061, p= 0.0004).

### Light-dark box test

Light-dark box data are summarized in Figure 3. ANOVA of time spent in the dark revealed significant effects of age (F_1, 35_ = 28.63, p < 0.0001), sex (F_1, 35_ = 8.101, p = 0.0074), and the interaction of age x sex (F_1, 35_ = 8.175, p = 0.0071). Sidak’s multiple comparison test revealed significantly more time in the dark in young vs aged NS females (p = 0.0022), young vs aged PRS females (p = 0.0047), and young PRS females vs young PRS males (p = 0.0022).

**Figure 3:**
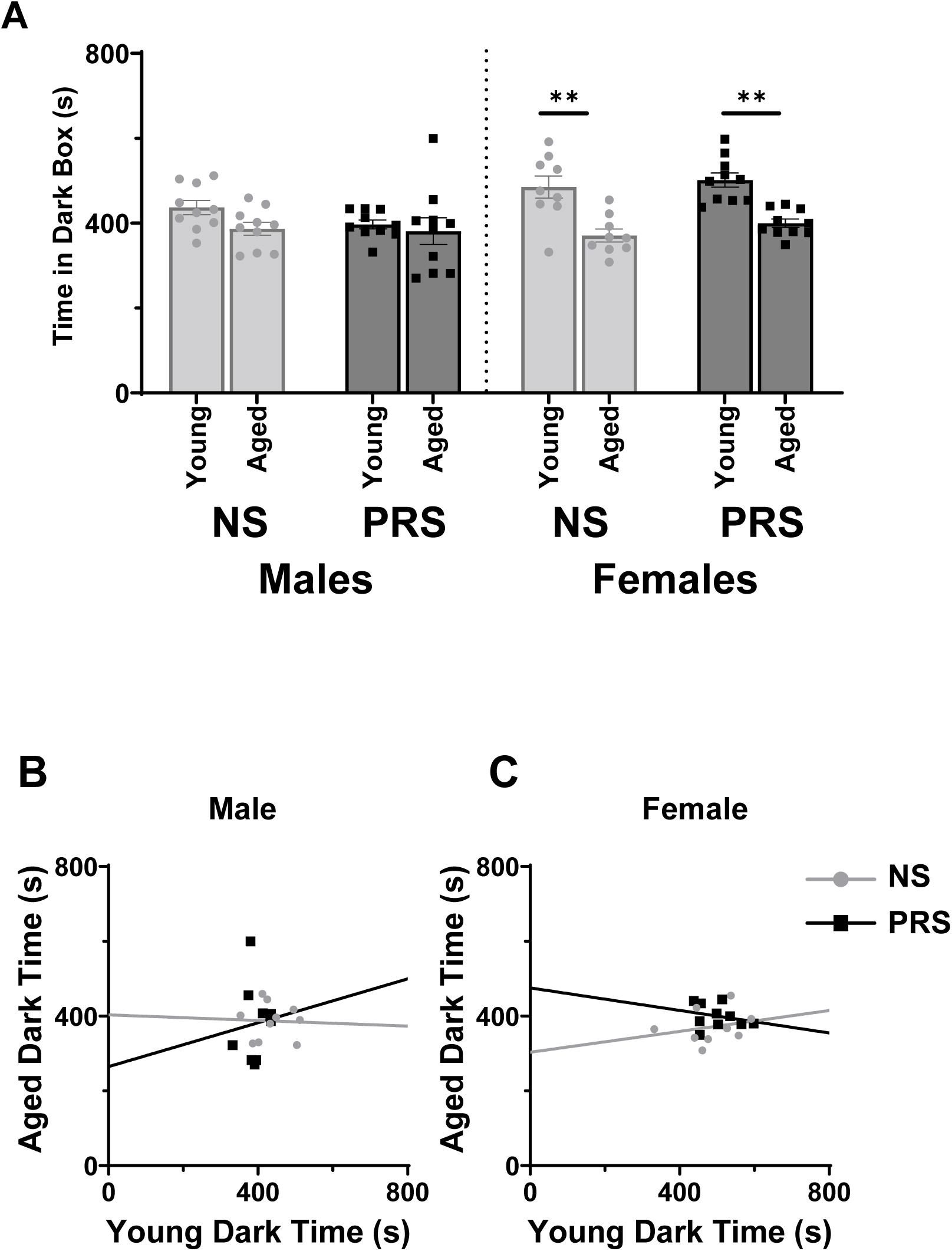
Light-dark box test of risk tolerance in mice. **A.** Comparisons of time spent in the dark side of the box during the 10-minute testing session between NS and PRS mice. Measures were taken at young adult and aged adult stages in the same mice. **B, C.** The correlations of light-dark box performance between mice at young and aged time points, for males and females, respectively. Multivariate ANOVA analysis resulted in a significant interaction between age and sex (F_1, 35_ = 8.175, p = 0.0071), but not age and stress. * p < 0.05, ** p < 0.01, *** p < 0.001, **** p < 0.0001

Simple linear regression of the time spent in the dark chamber between young and aged subjects revealed no significant deviations from 0 with R^2^ values all below 0.1.

### Morris Water Maze – cued learning test

ANOVA was performed to test a model including day x sex x stress variables. The ANOVA found a significant effect of day (F_1.193, 42.94_ = 65.88, p < 0.0001), but no other significant effects. As can be seen in the Morris water maze cued learning chart (Supplementary Fig. 2) all cohorts improved their latency to platform by day 3. Day 3 mean latency to platforms were 4.910, 8245, 6.023, and 6.215 for NS male, PRS male, NS female, and PRS female respectively. The cued learning was not performed in the aged cohort due to the test primarily being used as a control to determine whether mice were capable of performing the Morris water maze task.

### Morris Water Maze – spatial acquisition test

In each cohort during Morris water maze spatial acquisition, the mice were able to successfully learn the task (Fig. 4).

**Figure 4:**
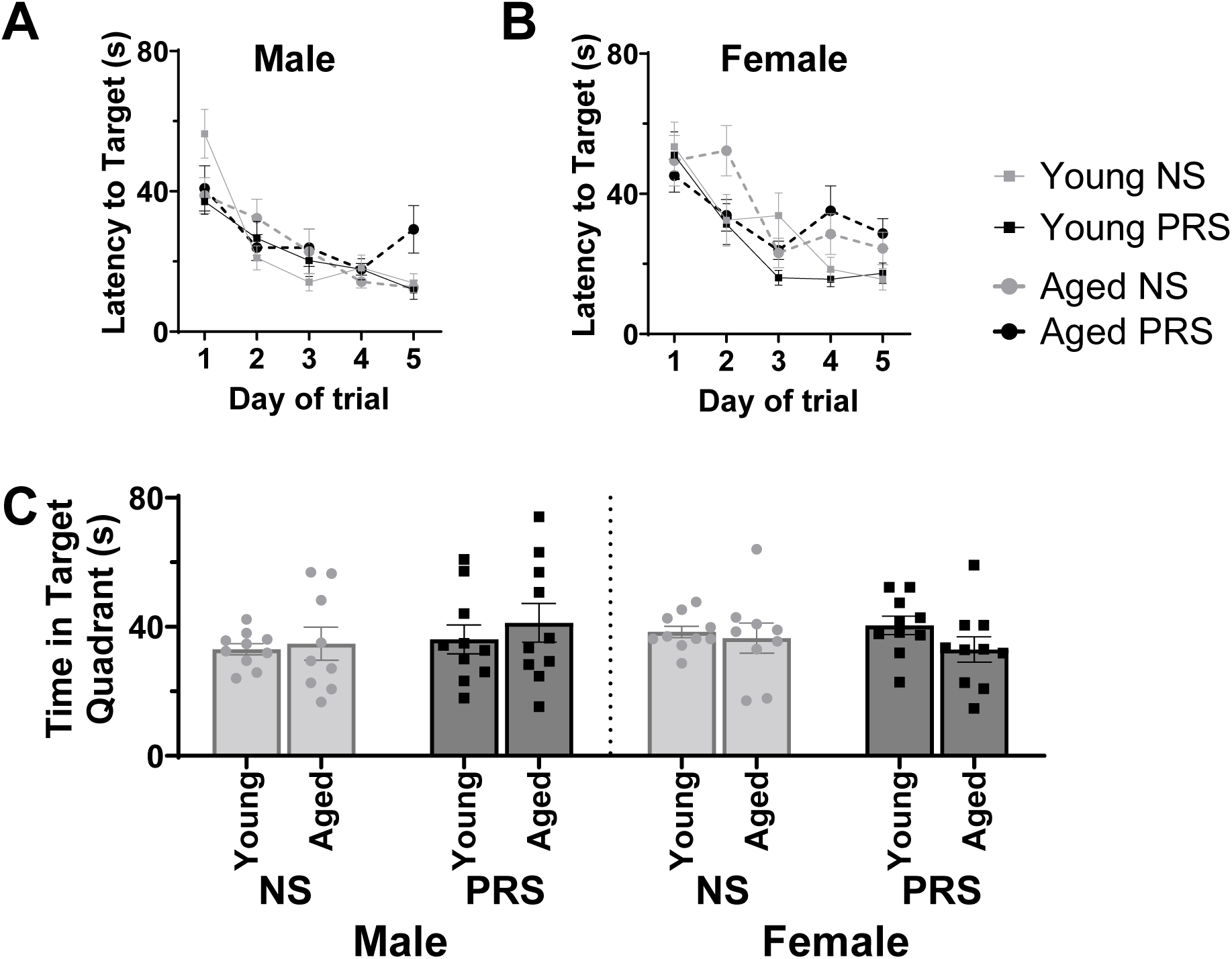
Morris Water Maze Spatial Acquisition. **A.** Male and **B.** female NS and PRS mice were tested for spatial acquisition which was performed for 5 days, with the maximum allotted time for each trial being 90 seconds before the subject was guided to the platform. Spatial acquisition was measured by the latency of time taken to reach the hidden platform, averaged by 4 trials each day from the different quadrants. **C.** Spatial acquisition memory was measured in a probe trial. The platform was removed prior to starting the probe trial and all mice were allowed to swim freely for 90 seconds. Memory was considered to be indicated by the time spent in the location of the platform was in during the tests of spatial acquisition learning. * p < 0.05, ** p < 0.01, *** p < 0.001, **** p < 0.0001

For latency to platform, a mixed-effects analysis of day x age x stress was performed for males and females separately rather than ANOVA due to one subject from both males and female groups dropping out during the Morris water maze trials. The mixed effects analysis was performed again to compare males and females in the probe trial for spatial acquisition.

For male latency to platform in spatial acquisition, three-way mixed effects analysis found a significant effect of day (F_4, 72_ = 26.28, p < 0.0001) because all groups of mice were learning the task. There was a significant interaction of day x age x stress because PRS males took slightly longer on a few days to find the hidden platform (F_4, 67_ = 3.073, p = 0.0219). For females, there was a significant effect of day (F_4, 72_ = 22.07, p < 0.0001), age (F_1, 18_ = 4.768, p = 0.0425), and a significant interaction between day x age (F_4, 67_ = 3.219, p = 0.0177). Young PRS females performed better than other female groups.

In the spatial acquisition probe trial, time spent in the target quadrant was a measure of spatial memory. Using a mixed-effects analysis, we found no significant effects of age, sex or stress on spatial memory. Furthermore, linear regression analysis showed that all groups except the NS males showed a positive correlation between the young vs. aged measurements of time in the target quadrant, but these differences were not statistically significant (Fig. 6A+B). Young adult PRS mice had a greater level of organized tracking behavior relative to NS mice (Fig. 7B). We observed trends for sex differences in spatial acquisition in aged mice, with a slightly higher level of organized tracking behavior in males relative to females (Fig. 7B).

### Morris water maze – spatial reversal test

Similar analyses were performed for the spatial reversal task as were done for the spatial acquisition task previously discussed. The Morris water maze spatial reversal test latency to platform times continuously improved in both males and females (Fig. 5A+B). All groups seemed to reach a similar average around 20 seconds by day 5 of testing. The time in the target quadrant for the reversal probe trial showed that male mice consistently spent around 40 seconds in the target quadrant regardless of stress or age (Fig. 5A). Female NS performed poorly when aged compared to measures when younger or in comparison with the male groups (Fig. 5B).

**Figure 5:**
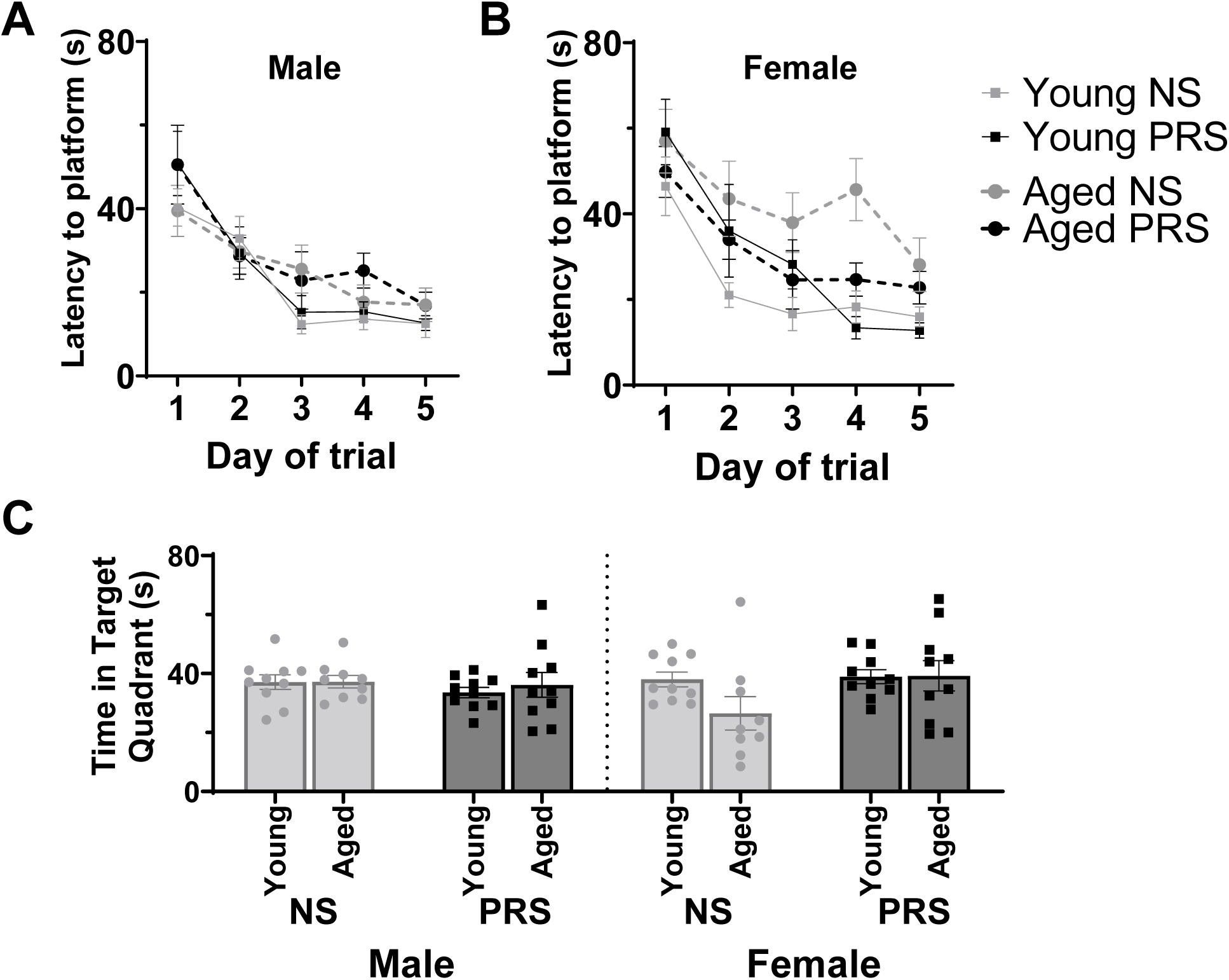
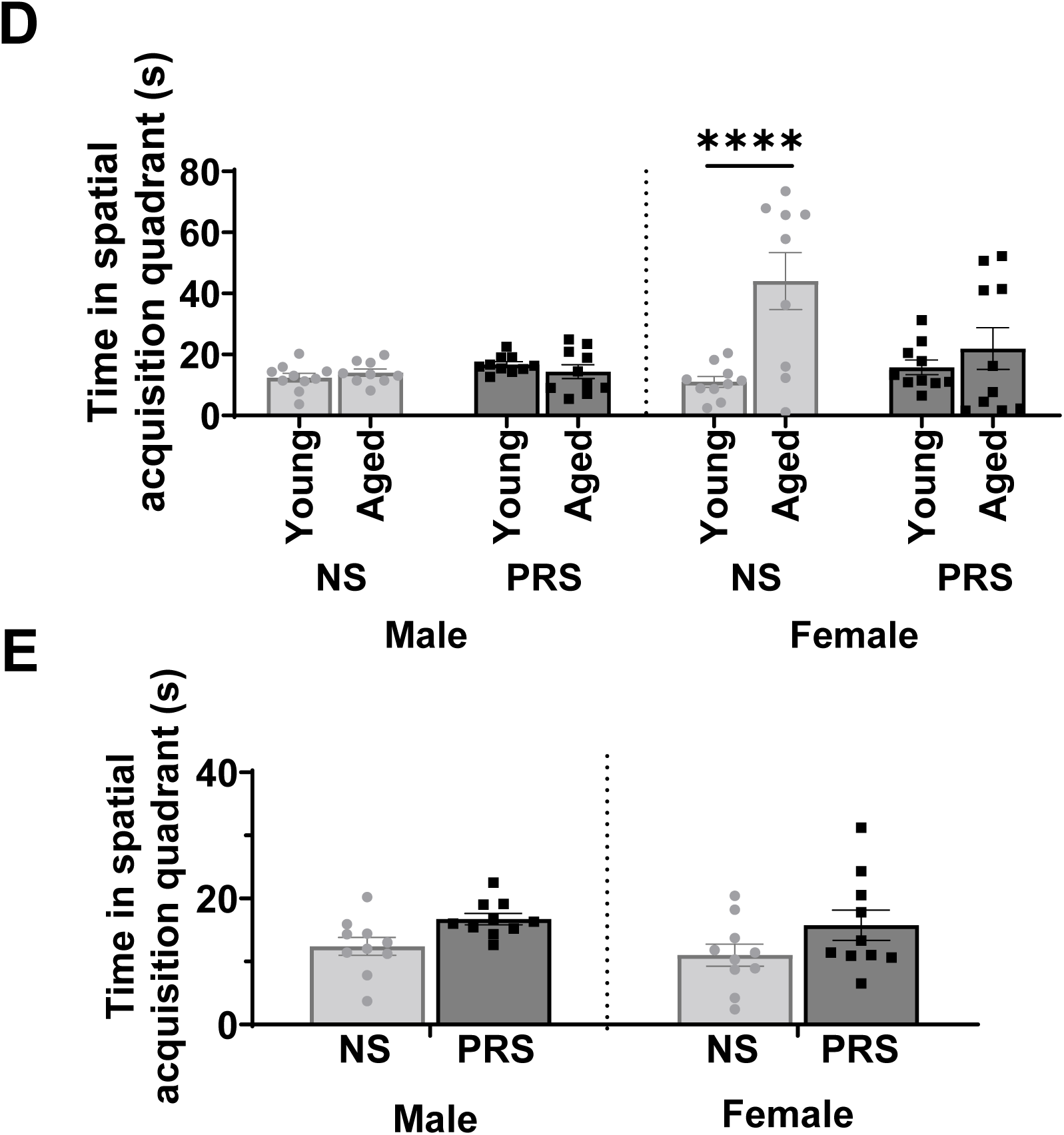
Morris Water Maze Spatial Reversal. After spatial acquisition testing, the hidden platform was moved to a new location. This test attempts to measure cognitive flexibility by measuring the latency to find the hidden platform in its new location. As with the spatial acquisition test, each trial day was averaged across four different starting positions. The maximum latency to platform was 90 seconds before the subjects were guided to the platform. **A,** Male groups did not differ in spatial reversal learning. **B.** Female aged NS subjects performed worse relative to measures in young adults. **C.** Spatial reversal probe trial was conducted after the hidden platform was removed. Time spent in the quadrant where the hidden platform was located during spatial reversal learning tests is a measure of the spatial reversal memory. **D.** We measured the time spent in the quadrant where the spatial acquisition platform was located previously, during the spatial reversal probe trial to test whether subjects had difficulty in extinguishing remote memory. **E.** We tested remote memory in young adult ages separately, which indicated increased remote memory of PRS mice for the spatial acquisition platform location. * p < 0.05, ** p < 0.01, *** p < 0.001, **** p < 0.0001.

In a mixed-effects analysis of male latency to platform during the spatial reversal task, there was a significant effect of day (F_4, 72_ = 32.22, p < 0.0001), but no significant effects of age or stress. For the females, there was a significant effect of day (F_4, 72_ = 27.53, p < 0.0001), age (F_0.6838, 12.31_ = 6.43, p = 0.0346), and a significant interaction between age and stress (F_1, 18_ = 4.924, p = 0.0396) as seen in Fig. 6D.

**Figure 6:**
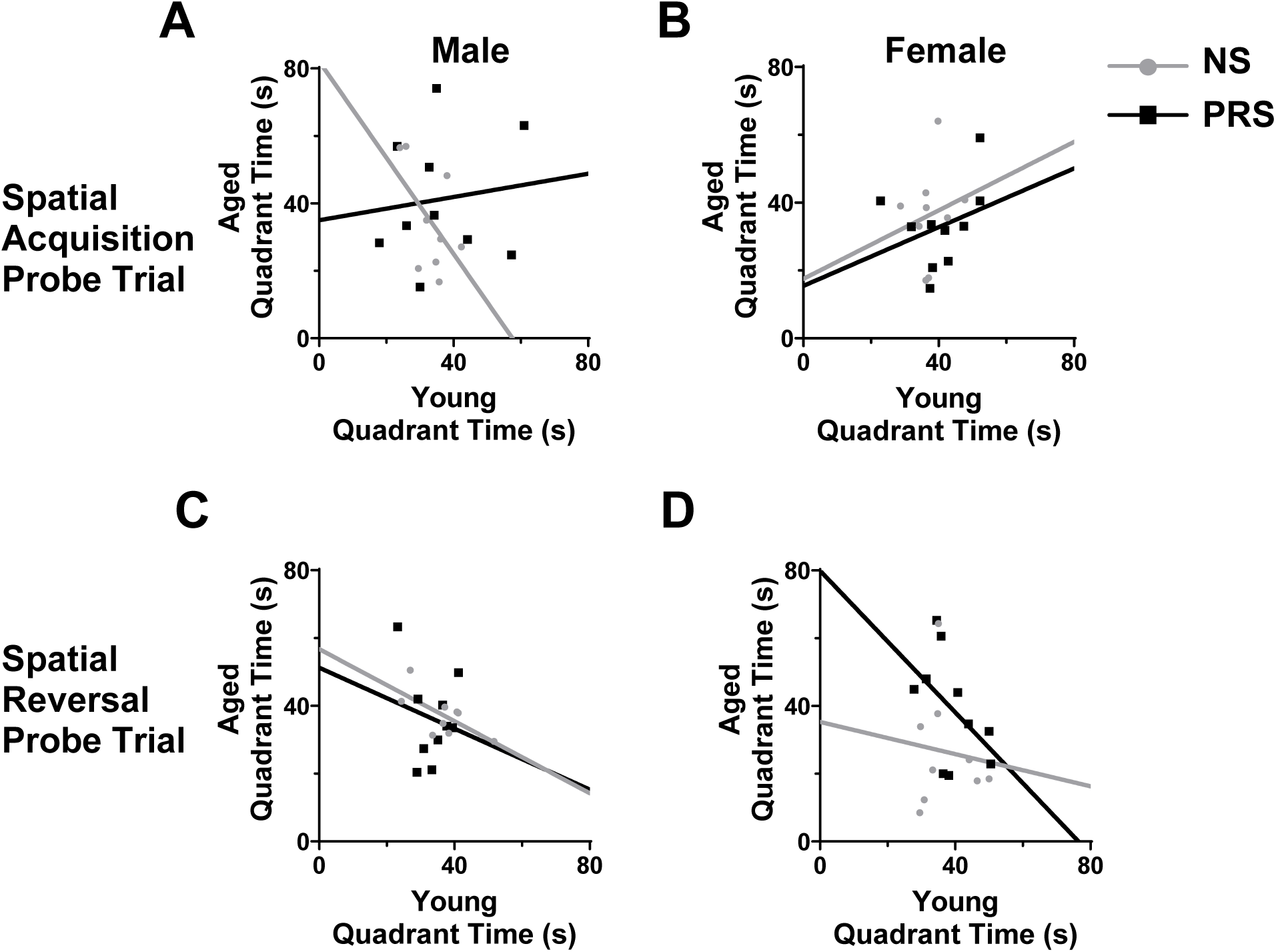
Morris Water Maze Probe Trials: Correlations between measures in young vs. old adult ages. A, B. Spatial acquisition probe trial measures in males and females at young and aged stages were correlated. Multivariate analysis suggested no significant interaction between age, sex, or stress. **C, D.** The time in the target quadrant for the spatial reversal task probe trial correlated between young and aged in males and females respectively. Multivariate analysis of the Morris water maze spatial reversal probe trial showed no significant interactions between age, sex, and stress.

In the spatial reversal probe trial, time spent in the target quadrant was not significantly associated with age, sex, or stress. We also looked at time spent in the original target quadrant from spatial acquisition to see if there were any issues with extinction of the initially acquired target location. In Figure 6D and 6E, time in the spatial acquisition target quadrant is shown. A mixed effects analysis of age x stress x sex showed significance of age (F_1,70_ = 10.40, p = 0.0019), Sex (F_1,70_ = 8.720, p = 0.0043), age x sex (F_1,70_ = 11.20, p = 0.0013), and age x stress (F_1,70_ = 6.690, p = 0.0118). Due to concerns over bias to one platform in the aged cohort, we did a separate analysis of the measures in early adulthood using a two-way ANOVA of sex x stress, which showed a significant effect of stress (F_1, 36_ = 7.033, p = 0.0118). Simple linear regression of the spatial reversal probe trial time in target quadrant comparing young and aged revealed a significant correlation with age in NS males (R^2^ = 0.4492, p = 0.0482), but no significant correlations between measures in young and old ages in the other groups (PRS male, NS female, and PRS female) (Fig. 6C). Female mice exposed to PRS had the most disorganized tracking behavior in the maze during the reversal trial relative to NS mice, even though this group had the most organized tracking behavior during spatial acquisition (Fig. 7B).

**Figure 7.**
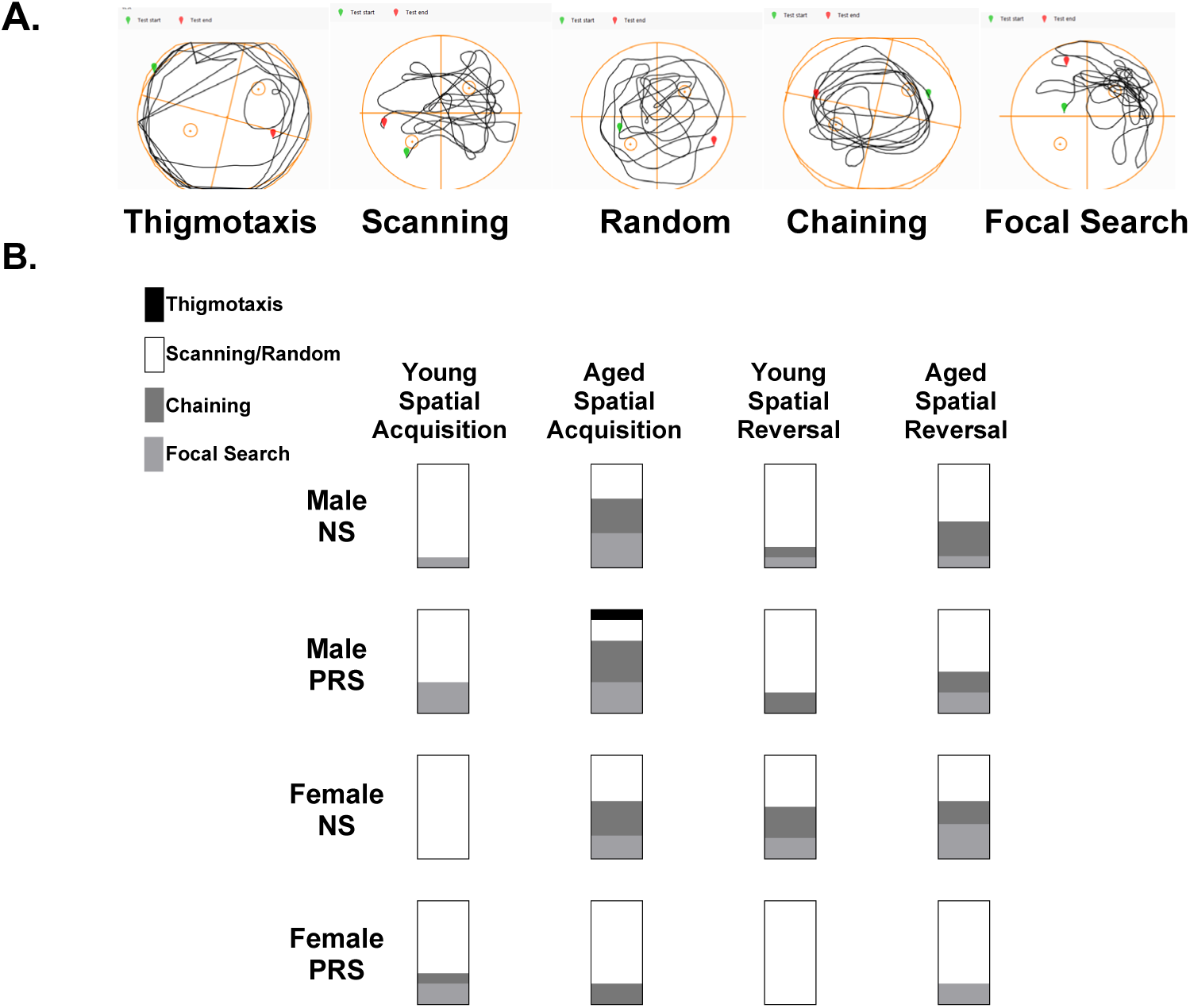
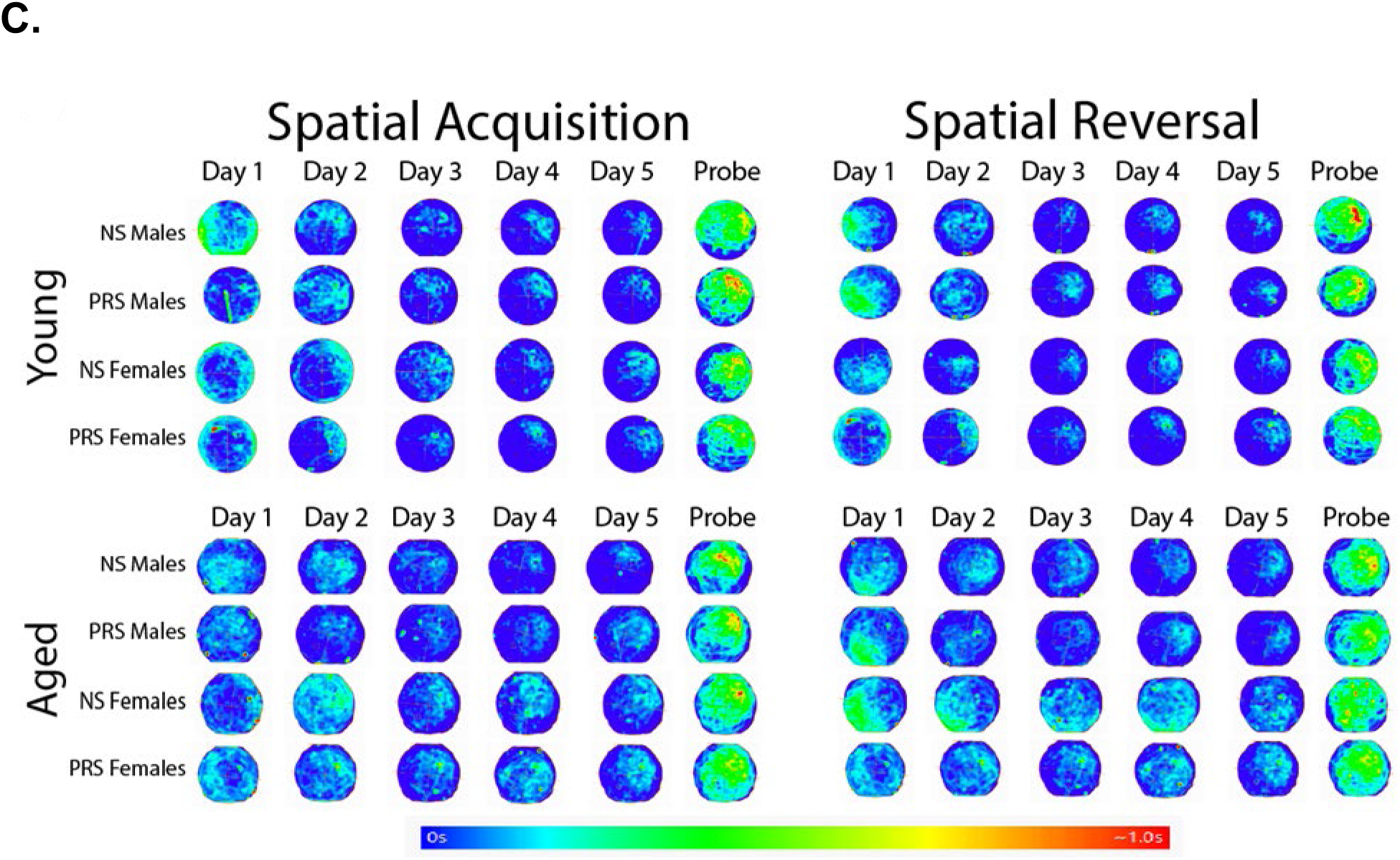
Track-plot categorization of Morris water maze probe trials. **A.** Summary images of trackplot categories of search strategies employed by mice in our experiments. The starting position of a test is indicated by a green marker and the location of the mouse and the end of the test is indicated by a red marker. Disorganized tracking behavior (thigmotaxis, scanning, random) and organized behaviors (chaining, focal search) are illustrated from representative plots in our mice. **B.** We identified significant differences between NS and PRS, ages of measurement and sexes in the spatial acquisition and spatial reversal when considering tracking behavior. The plot categories are summarized for each subgroup. Spatial acquisition trial (χ^2^=4.33 , df=1, p=0.038) performed in young adults because PRS mice had a greater level of organized tracking behavior relative to NS mice. However, the young PRS mice had a greater level of disorganized tracking behavior in the subsequent spatial reversal test (χ^2^=3.58, df=1, p=0.058). NS mice had more disorganized tracking behavior as young adults relative to their performance in the test at the later age, and relative to PRS mice at both ages (χ^2^= 9.29, df=1, p=0.0023). Aged mice had more organized tracking behavior in the spatial acquisition test than all younger groups, perhaps because they had repeated exposure to the maze (χ^2^=13.8 , df=1, p=0.0002). We observed trends for sex differences in spatial acquisition in aged mice, with a slightly higher level of organized tracking behavior in males(χ^2^=3.80, df=1, p=0.051). **C.** Heatmap images, generated from Any-Maze. Each map is an average of all subjects in each group (NS or PRS, male or female, young or aged) by day. Island locations were rotated to the northeast quadrant using Any-Maze; all goal locations are in the top right of each image. The intensity of red color is greatest where the average of all subjects reached 1 sec while the intensity of blue color is greatest where the average of all subjects was 0 secs. It should be noted that the aged female subjects had greatest variation in their paths leading to more dispersed areas of light blue through day 5. NS female aged subjects in the spatial reversal probe trial had the highest density at the previous location of the spatial acquisition island was located previously (the opposite quadrant) suggesting an increased memory of previous island locations.

### Social interaction and social memory

The percentage of time (out of the total 600 seconds) spent interacting with a stranger mouse relative to the empty cage was used as an index of social interaction or sociability. Also, a sociability index was calculated using entries in novel 2 cm surrounding zone divided by total entries into either of the 2 cm surrounding zones around each cage.

ANOVA between social v empty x sex x stress was used to analyze each of the parameters. There was a significant effect of social v. empty (F _1, 36_ = 175.1, p < 0.0001) but no other significant effects. *Post-hoc* analysis using Sidak’s multiple comparisons test revealed a significant difference between stranger and empty cups in each of NS male, PRS male, NS Female, and PRS female suggesting that the task was completed successfully. However, there were no significant differences when comparing the stranger interaction percentage times between each group. For sociability index, there was again a significant effect of stranger v empty (F _1, 36_ = 147.6, p < 0.0001) indicating that the mice showed a significant preference for the stranger. *Post-hoc* analysis using Sidak’s multiple comparisons test found similar significant differences between stranger and empty cups, but no other significant differences between groups.

For **social memory**, ANOVA comparing novel /familiar x sex x stress revealed a significant effect of novel/ familiar (F_1, 36_=15.36, p=0.0004), but no other significant effects. *Post-hoc* analysis using Sidak’s multiple comparisons test revealed no significant differences between any group in novel and familiar mice percentages. For sociability index in social memory, ANOVA revealed a significant effect of novel/ familiar, but no significant effects of sex or stress. *Post-hoc* analysis using Sidak’s multiple comparison test again showed there were non-significant differences between each group’s novel and familiar sociability indexes.

## Discussion

We sought to determine the effects of prenatal stress on cognitive behavior associated with the cortico-hippocampal circuit in C57BL/6J mice, which is the most commonly used inbred mouse strain in neurobiological studies due to its resilience to stress in animal facilities. This is the first detailed analysis of the effects of prenatal stress on spatial memory, including reversal learning in both sexes, in young adult and older adult mice. We discuss our findings in detail below.

### Prenatal stress alters spatial learning and memory

In the *<u>spatial acquisition test,</u>* we considered increased time taken (latency) to find the hidden platform in the Morris water maze as a negative indicator of spatial memory (Fig.4). Aged non-stressed females had the worst spatial acquisition latency relative to other groups. Age and stress had no effect on spatial acquisition latency in the male groups. Males and females performed similarly in the spatial acquisition probe trial. This is in contrast to findings in prenatally stressed mice by Mueller and Bale (2008), using the modified Barnes’ maze, and Clarke et al. (2019), using the Morris water maze. Both groups reported increased latency to escape and signs of decreased cognition in C57BL/6J mice. The reason for these differing findings may be due to different timing of prenatal stress exposure. Similar to our findings, Benoit et al. (2015) found no effect of prenatal stress on latency in the platform trials, but they did find a significant effect of prenatal stress on platform crossings during the probe trial. In addition to latency measures, the spatial navigation strategy used to find the hidden platform is also a measure of spatial cognition. Therefore, we tested the tracking strategies of mice in the Morris water maze as previously described (Cooke et al., 2019). We categorized these strategies as either ‘organized’ or ‘disorganized’ (Fig. 7). We observed that prenatal stress increased organized tracking strategies relative to other groups in young adult ages. However, the female prenatally stressed group performed worse than males and non-stressed females after aging. Search strategy analyses have revealed that the ventral hippocampus is involved in coarse spatial goal-directed search, that adult neurogenesis promotes spatially precise search, and that spatially accurate search is reduced in animal models that include hippocampal pathology (Cooke et al., 2019).

The effects of prenatal stress on *<u>spatial reversal</u>* has not been tested previously. We detected a significant effect of age and prenatal stress on reversal learning. Aged non-stressed females had reduced reversal learning compared to the other groups . Young and aged prenatally stressed females performed similarly, while the differential between measures in female non stressed mice at young and old ages appears to be much greater. Spatial navigation strategy in the reversal test was more organized in young and aged non-stressed females than young prenatally stressed mice of both sexes. In contrast males in both stress groups had more organized spatial reversal tracking behavior when aged (Fig. 7). This may hint at a susceptibility to decline in spatial cognition with age that did not occur in prenatally stressed females. Reversal learning in the Morris water maze can be used as a test of cognitive flexibility (Ragozzino & Rozman, 2007). Cognitive flexibility has been shown to be disordered in various neuropsychological diseases (Sanders et al., 2008), (Lange et al., 2017), (Uddin, 2021).

Our results show that cognitive flexibility measured by the spatial reversal in the Morris water maze appears to be reduced by prenatal stress in young mice. This may be caused by prenatal stress’s effects on the hippocampus (Fnaskova et al., 2023) which is important for spatial reversal learning (Shah et al., 2019). Interestingly, prenatal stress may be a protective factor as female mice age. Decreased cognitive flexibility can be seen in the heat maps of spatial reversal in aged NS females (Fig. 7C). In both the acquisition and probe trials, the densities appear more spread out rather than focal in the target quadrant. In human studies there is a similar age related decline in cognitive flexibility tested with a virtual Morris water maze which is moderated by neuropsychological functioning, including anxiety and depression (Schoenfeld et al., 2014). Direct tests of *<u>remote memory</u>*, have shown reduced extinction of those memories in late gestation prenatal stress models (Negron-Oyarzo et al., 2015), To test for failure to extinguish remote memory, we analyzed the time spent in the target quadrant of the spatial acquisition trials during the spatial reversal trial to see if there was any difficulty extinguishing memory of the earlier trial. At the young adult age, PRS groups of both sexes spent more time in the initial target quadrant, indicating a reduced ability to extinguish the memory of the earlier target location. Analysis of the aged cohort showed much higher time in the initial target quadrant in the NS female aged group compared to the other groups. Although slightly detrimental in early adulthood, prenatal stress may provide some protective benefit to hippocampal function as female mice age. Furthermore, simple linear regression analysis between young and aged subjects revealed a moderate correlation between NS males at young and older ages, while the other groups had no statistically significant correlations between their times at young and older ages, which may suggest that outcomes in NS males have more consistent hippocampal function through the lifespan compared to the PRS groups.

### Open field test for locomotor activity and risk aversion

*<u>Distance traveled</u>* revealed a significant effect of age and sex but not prenatal stress. Young males traveled a shorter total distance relative to older males and all female groups. When we considered *<u>center/periphery ratio</u>*, we also found no differences in any of the C57BL6/J groups tested. Similarly, Mueller and Bale (2008), found no stress-associated difference in the open field behavior when they tested the C57BL/6J subjects albeit using different prenatal stress exposure conditions. Clarke et al. (2019) reported hyperactivity in open field in prenatally stressed C57BL/6J mice. In contrast with conflicting findings in C57BL/6J, there are consistent findings of hyperactivity after Swiss Webster mice are exposed to prenatal stress (Dong et al., 2015; Matrisciano et al., 2013; Matrisciano et al., 2012; Moura et al., 2020).

Young males spent more *<u>time immobile</u>* in the open field, but this effect was attenuated with age and not altered by prenatal stress. Alternative measures of risk aversion or activity such as the zero maze, Behan et al. (2011) and Sierksma et al. (2013) also found no significant effects of prenatal stress. Moreover, we detected no difference between the stress groups at any age in the light-dark box test, but there were significant effects of age and sex. These data suggested that young females may be more risk averse than young males and become less risk averse as they age. Clarke et al. (2019) reported similar findings in C57BL/6J mice. Therefore the data indicate that prenatal stress does not reduce risk tolerance (previously referred to as ‘anxiety-like behavior) in the open field or light/dark box in C57BL/6J mice in young or older adults of either sex.

### Social Interaction and Social Memory Tests

In the *<u>social interaction test</u>*, all the mice spent significantly more time near the interaction cage than the empty cage, indicating the test was successful in measuring their preference for sociability. Dunn et al. (2024), using C57BL/6J, showed a decreased sociability index in prenatally stressed mice but we did not replicate those findings. In contrast, Clarke et al. (2019) found increased social interaction in prenatally stressed C57BL/6J mice. More consistent data between laboratories has been reported in Swiss webster mice, where prenatal stress decreases social interaction in male offspring (G. Bristow et al., 2021; Matrisciano et al., 2013; Matrisciano et al., 2012). The effect of prenatal stress on *<u>social memory</u>* has not been tested previously to our knowledge. Our data reveal no effect of prenatal stress on social novelty preference in C57BL/6J mice of both sexes and age groups (Fig. 8).

**Figure 8:**
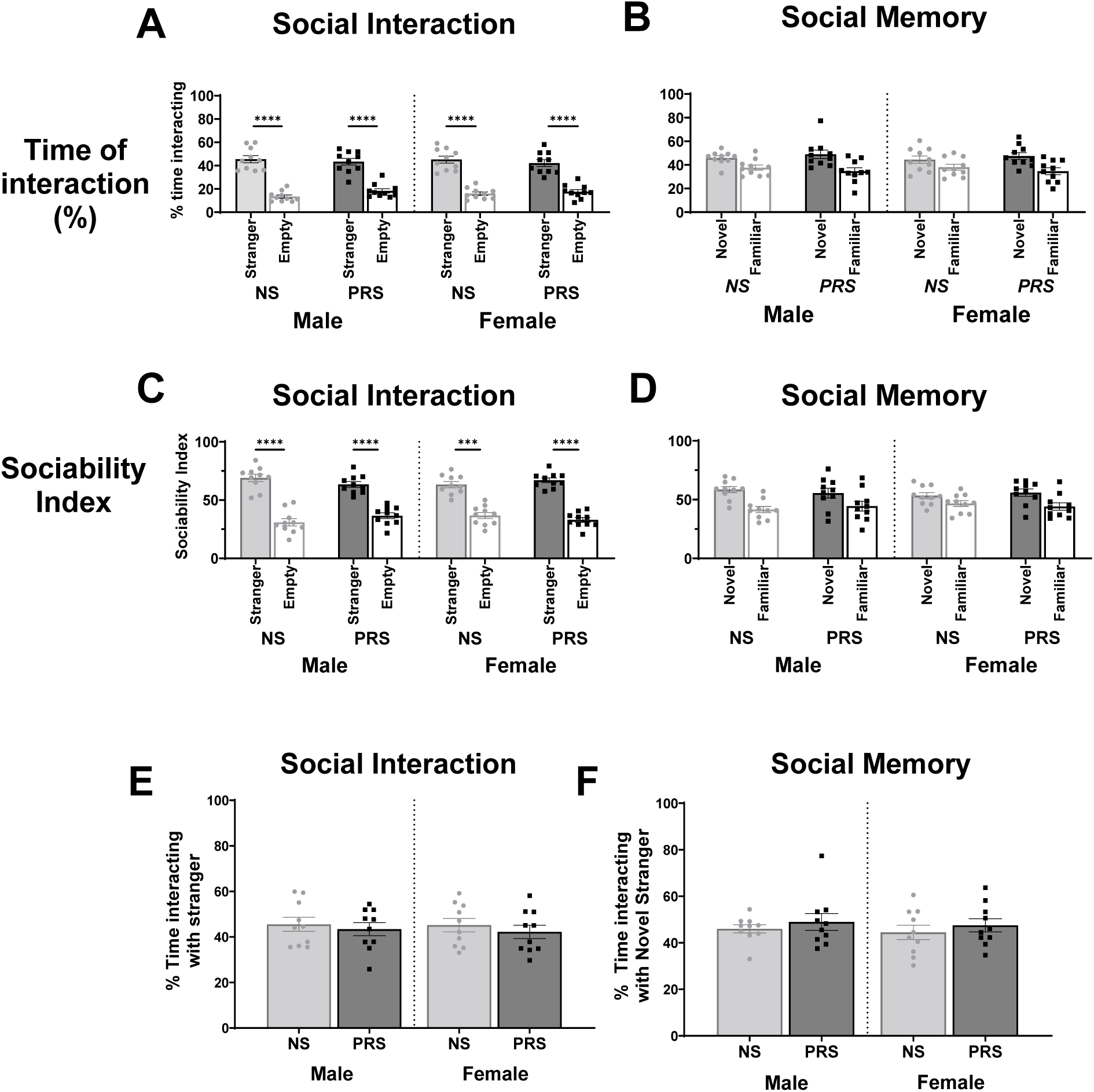
Social Interaction and social novelty comparisons. Graphs summarize social behavior measured by the percentage of total time interacting with each of the novel stranger cages. **A.** For social interaction, the white bars represent the time spent with the empty cage while the shaded bars represent time with the stranger mouse. **B.** For social memory, the white bars represent time spent with the familiar stranger while the shaded bars represent time spent with the novel stranger. **C, D.** The same coloring is applied to the sociability index, which was defined as entrances into the 2 cm surrounding circle divided by the total entries into <u>each</u> surrounding circle. **E.** Percentage time with stranger or **F.** novel stranger to compare the effects of stress in males and females. *** p < 0.001, **** p < 0.0001.

### Body weight

The total body weight for C57BL/6J male mice was reduced by prenatal stress. Dong et al. (2015) and Matrisciano et al. (2013) showed that Swiss Webster mice exposed to prenatal stress had significantly lower body weight at birth, but this difference diminished in early adulthood. The lower body weight at birth that we observed may have been attributable to the earlier birth time because PRS mice were born 8-12 hours earlier than NS mice. Mueller and Bale (2008), in did not report altered body weight in prenatally stressed C57BL/6J mice at 10-16 weeks of age.

### Differences between genetic backgrounds of mouse strains

We examined genetic differences that might increase resilience to prenatal stress in C57BL/6J mice relative to the more susceptible Swiss Webster strain from our previous study (Bristow et al., 2021). We summarize genetic differences between the strains in Supplementary Table 1. The Disrupted in Schizophrenia-1 (ΔDisc1) mutation in Swiss Webster mice is an obvious suspect for this strain’s relative susceptibility to prenatal stress. A *Disc1* translocation has been reported to co-segregate with schizophrenia in a Scottish family (Millar et al., 2000). A *Disc1* deletion reported in 129S6/SvEv mouse strains is associated with consistent working memory impairments when transferred to C57BL/6J mouse strains (Koike et al., 2006). This may be because the DISC1 protein plays a role in neurite outgrowth and cortical development (Koike et al., 2006).

The genetic basis of the relative resistance of the C57BL/6J strain to prenatal stress could be due to several genes that vary between the two mouse strains. . The *Ahr* gene which encodes the aryl hydrocarbon receptor has been associated with stress sensitivity (Madison et al., 2023). Another gene, with a better known neurobiological role that differs between C57BL/6J mice and Swiss Webster mice is the *Gabra2* gene. Specific Gabra2 haplotypes have been associated with addiction in human subjects (Enoch et al., 2010). RNA editing may also play a role in the pathology associated with prenatal stress (Bristow et al., 2021). Research directly comparing C57BL/6J to DBA/2J and BALB/cJ has shown increased serotonin turnover and variable RNA editing in each of the strains (Hackler et al., 2006).

We consider that our data supports the concept that C57BL/6J mice are more resilient to prenatal stress than other strains of mice (Crawley et al., 1997). Mueller and Bale (2007) and Behan et al. (2011) found limited evidence of an altered behavioral phenotype with late gestational stress. Dunn et al. (2024) found differences in social interaction sociability index and marble burying behavior. Like others, we do not observe these deficits in the C57BL/6J model with late prenatal restraint stress, and, thus, we consider this is primarily due to protective genomic variation in this mouse strain.

### Limitations of the study

Risk aversion or locomotor activity are potential confounds for performance in the Morris water maze. However, prenatal stress did not alter risk aversion behavior or locomotor activity in any group, therefore these confounds are unlikely. It is possible that the timing of prenatal stress may alter outcomes in offspring. It has been shown previously that late gestational stress in C57BL/6J models has limited negative neuropsychiatric outcomes in response to prenatal stress. More subtle deficits, like cognitive flexibility, may be tested further to investigate other possible neuropsychiatric issues associated with prenatal stress in the C57BL/6J model. Dunn et al. (2024) were able to show marble burying behavior is increased in the C57BL/6J strain in response to prenatal stress, and this may add to their findings of repetitive behaviors. A potential limiting factor could be unknown stressors within the animal housing facility.

Additionally, despite subjecting the dams to prenatal stress 3 times daily for eleven days, the dams may have received comparatively less stress through behavioral coping skills during the gestational restraint stress procedure such as maternal chewing behavior (Hisada et al., 2023). However, we have shown that prenatal stress impacts the behavioral outcomes in Swiss Webster mice, indicating that this prenatal stress exposure is successful in another strain of mouse. Lastly, neither estrogen nor testosterone cycles were standardized through the experiment so some female and male behaviors may have been affected by hormonal factors. However, Zhao et al. (2021) reported that estrus cycling in C57BL/6J had minimal impact in behavioral testing data.

### Conclusions

In summary, data reveal that male but not female mice had reduced body weight throughout the lifespan after exposure to prenatal stress. Young prenatally stressed adult females showed greater organization of tracking behavior in spatial acquisition tests, but prenatal stress improved reversal learning in aged females. Therefore the only deficit caused by prenatal stress that we observed in the C57BL/6J strain was in spatial reversal learning and memory, which is related to cognitive flexibility. Overall results indicate that prenatal stress may protect females from detrimental effects of age to the hippocampus. In conclusion, our data support the finding that C57BL/6J are a relatively resilient strain of mouse that may be useful for investigating the differences between resilience and susceptibility to stress. Future studies of prenatal stress in this mouse strain should focus on the identification of molecular pathways that underlie resilience to prenatal stress, to identify novel targets for drug development.

---

**Supplementary Figure 1:**
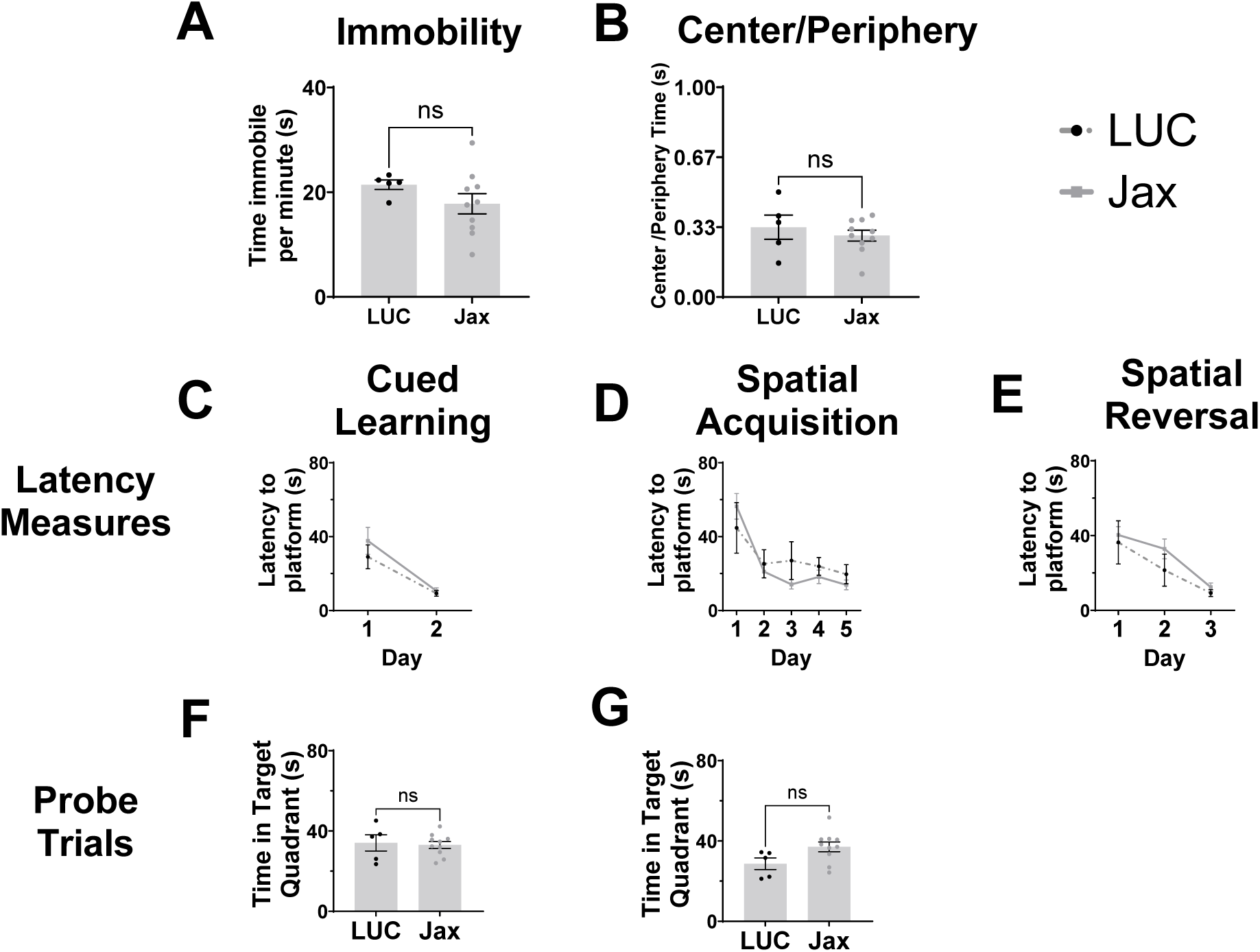
Comparison of male control subgroups. Litters bred at LUC were supplemented with male control mice purchased from Jackson Laboratories several weeks before the behavioral analyses. We compared behavioral measures between these control groups and find no significant differences. **A, B.** Open-field data when comparing the control subjects bred in house against the Jax-mice. **C-E.** Cued learning, spatial acquisition, and spatial reversal trials, respectively, did not differ between the control groups. **F. G.** Probe trials for spatial acquisition and spatial reversal, respectively, show no significant difference between the control groups. * p < 0.05, ** p < 0.01, *** p < 0.001, **** p < 0.0001

**Supplementary Figure 2:**
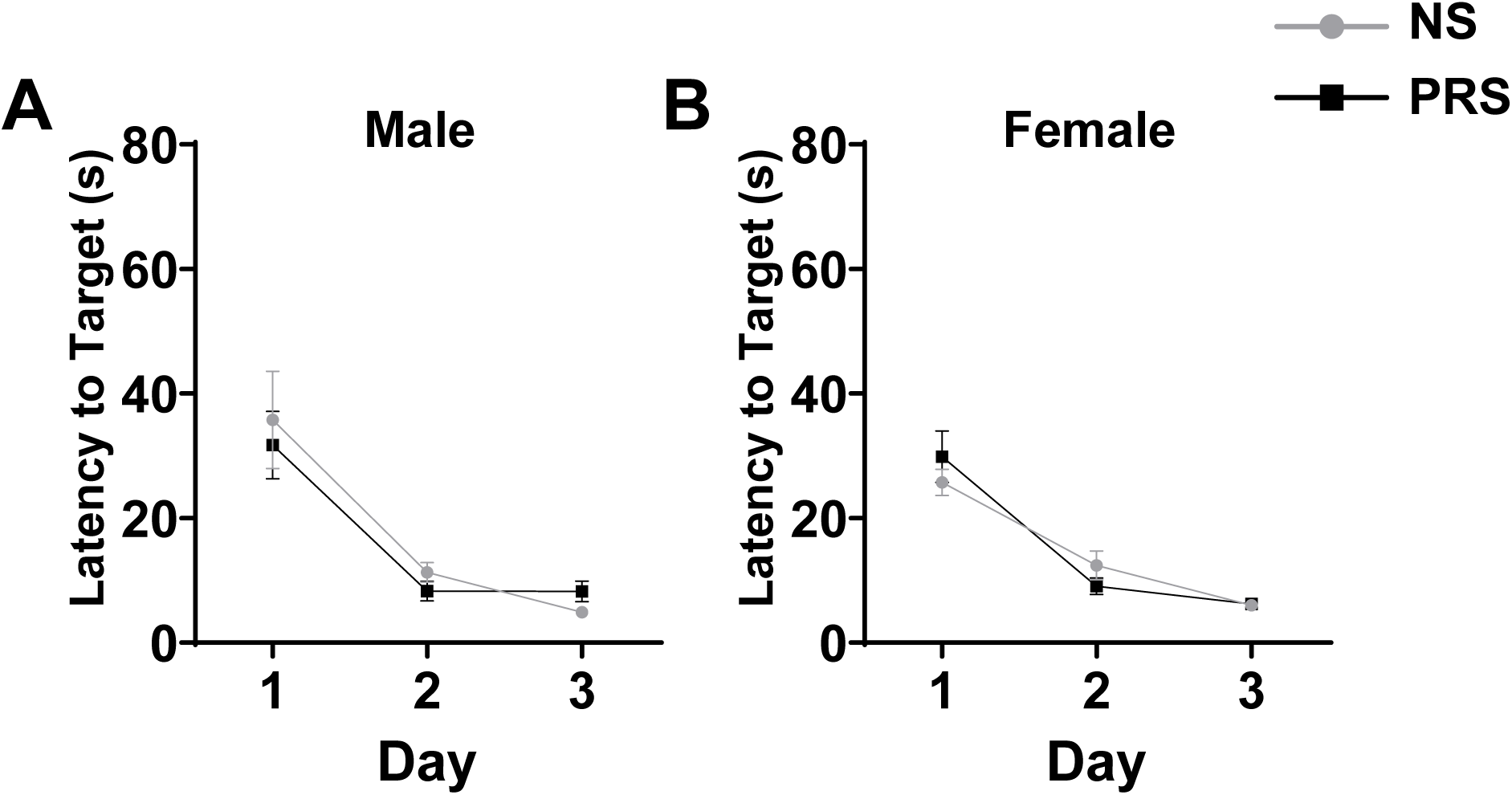
Morris Water Maze, Cued Learning. **A, B.** Cued learning data in males and females respectively. Each subject had 4 trials per day for 3 days in the maze. The platform was signposted using a yellow tennis ball.

**Supplementary Table 1:**
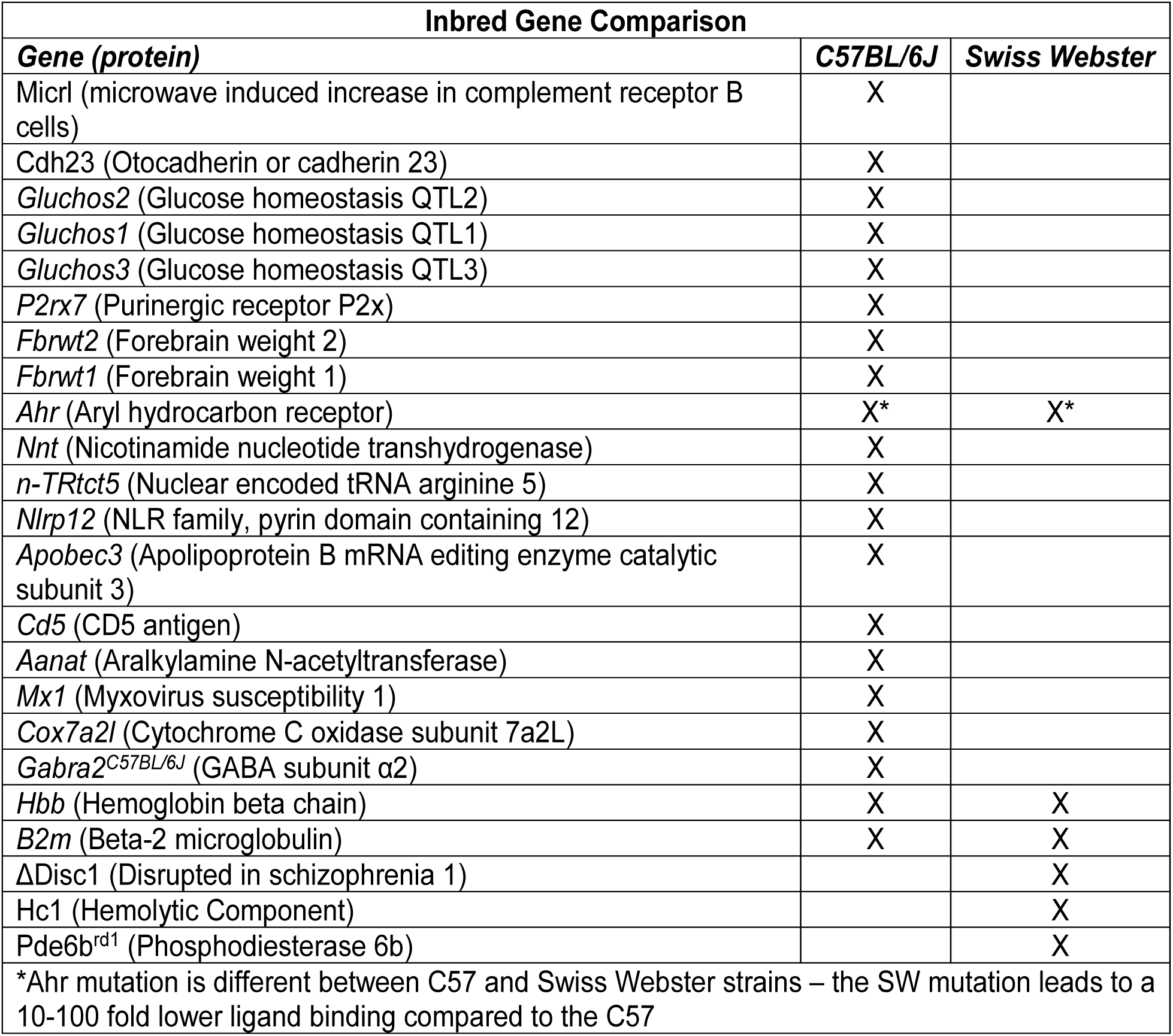
Inbred genetic comparisons. – made using Mouse Genome Informatics (www.informatics.jax.org); in-depth genetic information researched through GeneCards (www.genecards.org). X marks the presence of the genetic mutation within each inbred strain.

