## Supplementary Table 1, Supplementary Figure 1, Supplementary Figure 2 for "PRENATAL STRESS MODIFIES SPATIAL COGNITION IN MICE: EFFECTS OF AGE AND SEX"

**Supplementary Figure 1: Comparison of male control subgroups.**

Litters bred at LUC were supplemented with male control mice purchased from Jackson Laboratories several weeks before the behavioral analyses. We compared behavioral measures between these control groups and find no significant differences.

**A, B.** Open-field data when comparing the control subjects bred in house against the Jax-mice.

**C-E.** Cued learning, spatial acquisition, and spatial reversal trials, respectively, did not differ between the control groups.

**F. G**. Probe trials for spatial acquisition and spatial reversal, respectively, show no significant difference between the control groups. * $p < 0.05$, ** $p < 0.01$, *** $p < 0.001$, **** $p < 0.0001$

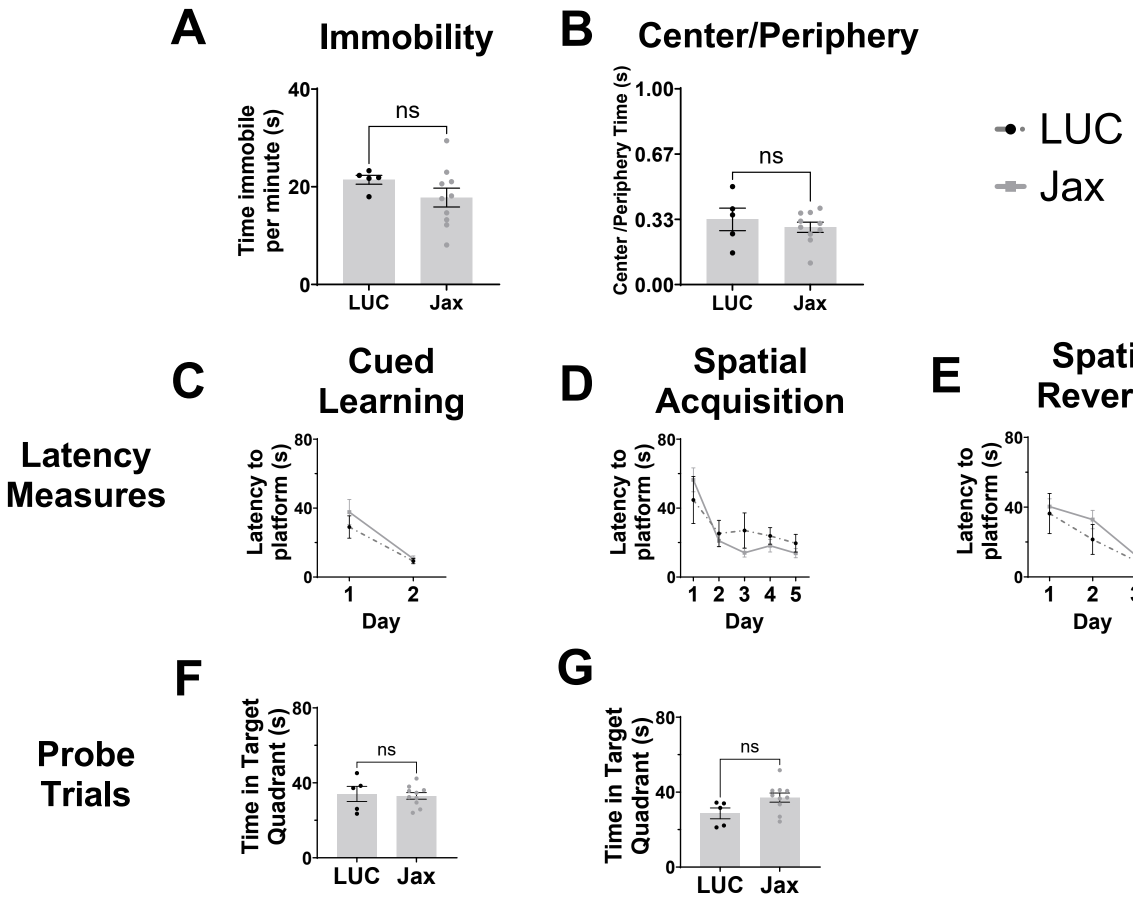

**Supplementary Figure 2: Morris Water Maze, Cued Learning**.
**A, B.** Cued learning data in males and females respectively. Each subject had 4 trials per day for 3 days in the maze. The platform was signposted using a yellow tennis ball.

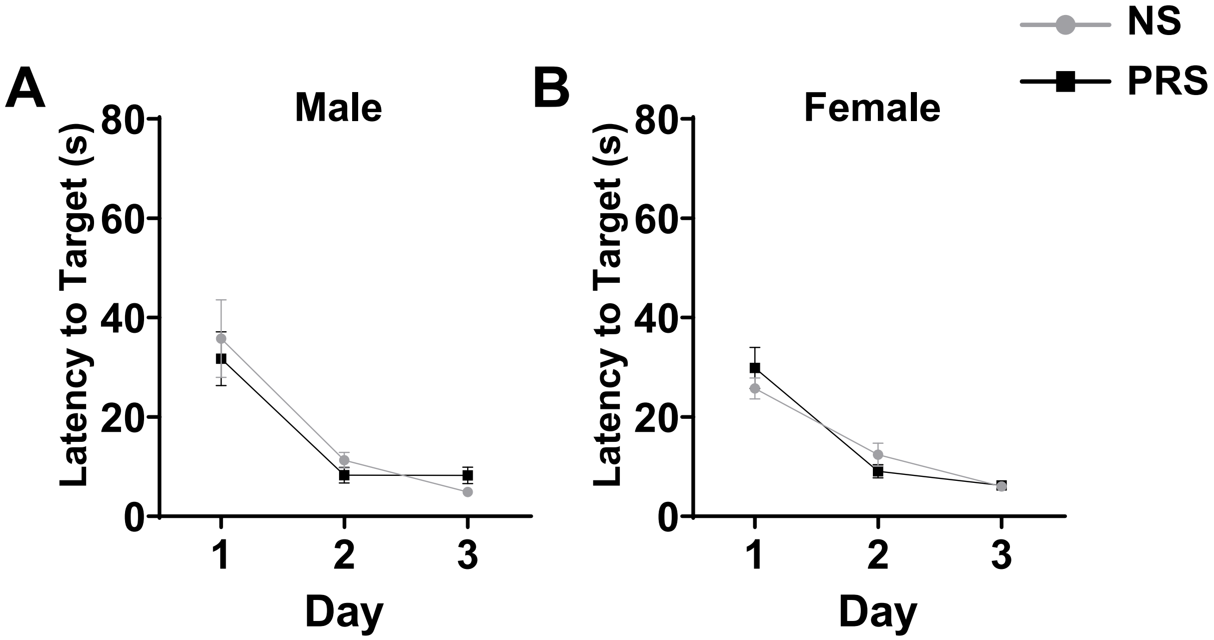

**Supplementary Table 1: Inbred genetic comparisons** – made using Mouse Genome Informatics (www.informatics.jax.org); in-depth genetic information researched through GeneCards (www.genecards.org). X marks the presence of the genetic mutation within each inbred strain.

| **Inbred Gene Comparison** | | |
|---|---|---|
| ***Gene (protein)*** | ***C57BL/6J*** | ***Swiss Webster*** |
| Micrl (microwave induced increase in complement receptor B cells) | X | |
| Cdh23 (Otocadherin or cadherin 23) | X | |
| *Gluchos2* (Glucose homeostasis QTL2) | X | |
| *Gluchos1* (Glucose homeostasis QTL1) | X | |
| *Gluchos3* (Glucose homeostasis QTL3) | X | |
| *P2rx7* (Purinergic receptor P2x) | X | |
| *Fbrwt2* (Forebrain weight 2) | X | |
| *Fbrwt1* (Forebrain weight 1) | X | |
| *Ahr* (Aryl hydrocarbon receptor) | X* | X* |
| *Nnt* (Nicotinamide nucleotide transhydrogenase) | X | |
| *n-TRtct5* (Nuclear encoded tRNA arginine 5) | X | |
| *Nlrp12* (NLR family, pyrin domain containing 12) | X | |
| *Apobec3* (Apolipoprotein B mRNA editing enzyme catalytic subunit 3) | X | |
| *Cd5* (CD5 antigen) | X | |
| *Aanat* (Aralkylamine N-acetyltransferase) | X | |
| *Mx1* (Myxovirus susceptibility 1) | X | |
| *Cox7a2l* (Cytochrome C oxidase subunit 7a2L) | X | |
| $Gabra2^{C57BL/6J}$ (GABA subunit α2) | X | |
| *Hbb* (Hemoglobin beta chain) | X | X |
| *B2m* (Beta-2 microglobulin) | X | X |
| ΔDisc1 (Disrupted in schizophrenia 1) | | X |
| Hc1 (Hemolytic Component) | | X |
| $Pde6b^{rd1}$ (Phosphodiesterase 6b) | | X |
| *Ahr mutation is different between C57 and Swiss Webster strains – the SW mutation leads to a 10-100 fold lower ligand binding compared to the C57 | | |
